# Perinatal maternal undernutrition in baboons modulates hepatic mitochondrial function but not metabolites in aging offspring

**DOI:** 10.1101/2024.05.02.592246

**Authors:** Daniel A. Adekunbi, Bowen Yang, Hillary F. Huber, Angelica M. Riojas, Alexander J. Moody, Cun Li, Michael Olivier, Peter W. Nathanielsz, Geoffery D. Clarke, Laura A. Cox, Adam B. Salmon

**Author notes:** **Corresponding author:** Dr. Adam Salmon, Barshop Institute for Longevity and Aging Studies, The University of Texas Health Science Center at San Antonio, 7703 Floyd Curl Dr, San Antonio TX, 78229. Texas, USA.

## Abstract

We previously demonstrated in baboons that maternal undernutrition (MUN), achieved by 70 % of control nutrition, impairs fetal liver function, but long-term changes associated with aging in this model remain unexplored. Here, we assessed clinical phenotypes of liver function, mitochondrial bioenergetics, and protein abundance in adult male and female baboons exposed to MUN during pregnancy and lactation and their control counterparts. Plasma liver enzymes were assessed enzymatically. Liver glycogen, choline, and lipid concentrations were quantified by magnetic resonance spectroscopy. Mitochondrial respiration in primary hepatocytes under standard culture conditions and in response to metabolic (1 mM glucose) and oxidative (100 µM H_2_O_2_) stress were assessed with Seahorse XFe96. Hepatocyte mitochondrial membrane potential (MMP) and protein abundance were determined by tetramethylrhodamine ethyl ester staining and immunoblotting, respectively. Liver enzymes and metabolite concentrations were largely unaffected by MUN, except for higher aspartate aminotransferase levels in MUN offspring when male and female data were combined. Oxygen consumption rate, extracellular acidification rate, and MMP were significantly higher in male MUN offspring relative to control animals under standard culture. However, in females, cellular respiration was similar in control and MUN offspring. In response to low glucose challenge, only control male hepatocytes were resistant to low glucose-stimulated increase in basal and ATP-linked respiration. H_2_O_2_ did not affect hepatocyte mitochondrial respiration. Protein markers of mitochondrial respiratory chain subunits, biogenesis, dynamics, and antioxidant enzymes were unchanged. Male-specific increases in mitochondrial bioenergetics in MUN offspring may be associated with increased energy demand in these animals. The similarity in systemic liver parameters suggests that changes in hepatocyte bioenergetics capacity precede detectable circulatory hepatic defects in MUN offspring and that the mitochondria may be an orchestrator of liver programming outcome.

## 1.0 Introduction

Suboptimal nutrition during the developmental period, from *in utero* through early life development, is common among individuals from disadvantaged populations. Nutritional perturbations during these critical periods can induce long-standing physiological changes that increase risk of developing chronic diseases, including metabolic diseases, later in life. In response to nutritional stress during pregnancy, the fetus initiates adaptive processes to ensure its survival albeit at the expense of optimal structural and functional development. The mechanisms by which early-life exposures and challenges increase susceptibility to adult-onset diseases are not entirely clear, but it is likely that adaptive process to ensure fetal survival likely come at the expense of optimal structural and functional development that persist throughout life^1,2^.

The liver has been shown to be keenly sensitive to *in utero* nutrient restriction^3^, and fetal liver size is reduced under this challenge in various species including rats, sheep, cattle, baboons, and humans among others^4–8^. We and others have also reported alterations in fetal liver metabolites and gene expression patterns due to maternal nutrient reduction^6,9–11^. Although several other animal models have shown association between maternal undernutrition (MUN) and disrupted metabolism in adulthood^12,13^, there is a need to study nonhuman primates due to their close phylogenetic relationship and similar physiology to humans^14^ to bridge the translational gap in developmental programming studies. We previously noted the emergence of insulin resistance in juvenile (3.5 years) offspring of baboons exposed to moderate MUN during pregnancy and lactation^15^. The liver undergoes several functional changes during the early postnatal period before achieving full maturation^16^ and developmental programming imprints persist throughout development and early adulthood of the offspring. Long-term changes associated with aging in this model remain unexplored.

Metabolic disorders are primarily driven by disruption in energy homeostasis, of which the mitochondria can play a key role. Mitochondria play multitude of roles in regulating energy balance including ATP production, generation of reactive oxygen species (ROS) and regulating cellular signaling pathways, and impairment of these systems contribute to metabolic dysfunction. Developmental programming of critical component systems such as the mitochondria has been proposed as a cellular mechanism by which maternal effects are propagated in the offspring given the developmental plasticity of the mitochondria and maternal imprints in the offspring mitochondrial genome^17^. Our recent study demonstrated that MUN impaired fetal mitochondrial structure and function, including alterations in mitochondrial cristae and bioenergetics in cardiac tissue^18^. Impairment in fetal mitochondrial function is linked to compromised metabolic health in postnatal life^19^, and cumulative damage to the mitochondria is suggested to trigger the onset of many age-related diseases^20^. In line with this, we have shown that MUN in baboons drives mitochondrial bioenergetic defects that persist even to adulthood in skin-derived fibroblasts^21^.

As a follow-up to our previous studies^10,11,13^, we utilized baboons with an average age of 15 years (approximate human equivalent; 60 years), representing animals in or transitioning to the late stage of life, to investigate the long-term impact of MUN on the liver at the functional and molecular level. In the present study, we assessed clinical phenotypes of liver function by analyzing plasma changes in liver enzymes to identify functional hepatic deficits related to MUN in the aging offspring. Additionally, we examined lipid accumulation and other metabolites in the liver using magnetic resonance spectroscopy. Considering the possibility that subcellular changes may present earlier than phenotypic alterations, that is, systemic outcomes are preceded by cellular changes, we assessed mitochondrial bioenergetics and cellular protein abundance in primary hepatocytes derived from these animals.

## 2.0 Methods

### 2.1. Animals

The Institutional Animal Care and Use Committee of Texas Biomedical Research Institute (TBRI) approved all procedures involving animals. The animal facilities at the Southwest National Primate Research Center (SNPRC), housed on TBRI campus are fully accredited by the Association for Assessment and Accreditation of Laboratory Animal Care International (AAALAC), and adheres to the guidelines of the National Institutes of Health (NIH) and the U.S. Department of Agriculture.

Details of animal husbandry and establishment of MNR model have been published previously by our group^22, 23^. Baboons (*Papio* sp.) were housed and maintained in a social environment and fed ad libitum with normal monkey diet. The welfare of the animals was enhanced by providing enrichments, such as toys, food treats, and music, which were offered daily under the supervision of the veterinary and behavioral staff at SNPRC.

The baboon colony used in this study were established more than 20 years ago. To develop the MUN cohort, age-matched females were randomly assigned prior to breeding to control or MUN group. Control mothers had *ad libitum* access to water and SNPRC biscuits (Purina Monkey Diet and Monkey Diet Jumbo, Purina LabDiets, St Louis, MO, USA) containing 12% energy from fat, 0.29% from glucose, 0.32% from fructose, and a metabolizable energy content of 3.07 kcal/g. MUN group were fed 70% of the feed eaten by the control females on a weight-adjusted basis from the time of diagnosis of pregnancy (∼30 days gestation) for the rest of pregnancy and through lactation. We have previously demonstrated that MUN leads to intrauterine growth restriction (IUGR) at term^24^. Both control and MUN offspring were weaned at 9 months and maintained on Purina Monkey diet through adulthood. Animals were euthanized at ages ranging between 13 and 18 years (approximate human equivalent, 50 and 70 years).

### 2.2. Magnetic resonance spectroscopy

All proton magnetic resonance spectroscopy (1H-MRS) experiments were conducted on a Siemens 3T system (Trio, Siemens Healthcare, Malvern, PA) with a transmitting body coil. The scans were carried out when the animals were approximately 14 years old. We used straps to minimize involuntary motion during the scanning protocols. MRI scans were performed while subjects were mechanically ventilated and under sedation according to the following protocol: After an overnight fast (12 h), each baboon was sedated with ketamine hydrochloride (10 mg/kg i.m.) before arrival at the MRI room. Endotracheal intubation was performed using disposable cuffed tubes (6.5 - 8.0 mm diameter) under direct laryngoscopic visualization. All animals were supported with 98 - 99.5% fraction of inspired oxygen (FiO_2_) by a pressure-controlled ventilator adjusted, as necessary, to keep the oxygen saturation >95%. The maintenance of anesthesia consisted of an inhaled isofluorane (0.5 - 1.5%) and oxygen mix.

Glycogen, choline, and lipid concentrations in the liver were quantified using single-voxel, spin echo localized 1H-MRS^25^. A voxel with a volume of 12×12×12 mm^3^ was placed in the right posterior love of the liver. The voxel was placed approximately 2 cm within Gleason’s capsule to avoid signal contamination from the visceral adipose compartment. Due to the amplitude of the water resonance, two spectra were collected for each subject: a water reference (TR = 2000 ms, TE = 30 ms, NSA = 8) and a water-saturated spectrum (TR = 2000 ms, TE = 30 ms, NSA = 16).

### 2.3. 1H-MRS data processing

All spectral peaks were fit using the non-linear least squares, an advanced method for the accurate, robust, and efficient spectral fitting algorithm (AMARES) in the Java-based magnetic resonance user interface software (jMRUI v5.2). The detailed process of analyzing 1H-MRS data has been previously described^25, 26^. Firstly, to reduce fitting residuals, MRS data are processed by fitting spectral peaks using the spectral-fitting algorithm in the MRS analysis software jMRUI. Spectra were corrected for phase offsets by applying a phase shift not exceeding ± 12 for either the reference or unsuppressed spectra. Secondly, if any residual water resonance was present for water-suppressed spectra, it was removed by applying the Hankel Lanczos singular value decomposition (HL-SVD) filter with no point maxima. The reference peak was assigned to the water peak (in unsuppressed spectra). Water-suppressed spectra were also filtered using apodization with a 3.5 Hz Gaussian. Water signals are generated from the water-unpressed spectrum. During the jMRUI analysis, starting values and prior knowledge estimates were applied according to previous publications^25, 27, 28^. Since there is an inherent signal loss at the point of data acquisition, glycogen, choline, and lipid signals were corrected by T2 relaxation^25, 27^.

### 2.4. Tissue Collection

At approximately 15 years of age, male and female adult baboons were tranquilized with ketamine hydrochloride (10 mg/kg i.m.) after an overnight fast. Three days prior to necropsy, morphometrics including body weight and length were determined to calculate body mass index and blood samples drawn through the femoral vein to obtain plasma for liver enzymes analyses. On the day of necropsy, tranquilized baboons were exsanguinated while still under general anesthesia as approved by the American Veterinary Medical Association. Following failure of reflex responses to skin pinch and eye touch stimulation, liver tissues were rapidly removed and weighed. Tissues were collected between 8.00-10.00 AM to minimize potential variation from circadian rhythm. Left and right liver lobes were separated apart, about 30 grams of each lobe was cut laterally closer to the caudal portion of the liver and received into an ice-chilled 1X HBBS for subsequent hepatocyte isolation. The remaining liver portion was immediately frozen in liquid nitrogen for other analyses as part of other investigations. All necropsies were performed by qualified and experienced veterinarians.

### 2.5. Plasma liver enzyme quantification

Markers of liver function including aspartate aminotransferase (AST), alkaline phosphatase (ALP), and alanine aminotransferase (ALT) were assessed in plasma samples using the Beckman Coulter UniCel DxC 800 Synchron Clinical System (Brea, CA, USA) along with their specific reagents from Beckman Coulter. AST, ALP, and ALT were measured using enzymatic rate method^29^.

### 2.6. Hepatocyte cultures

The two-step EGTA/collagenase perfusion technique^30, 31^ was adapted to isolate primary hepatocytes from baboon liver ex situ. Liver samples were processed within 1 h post collection to achieve viable hepatocytes. Two cannulae (16g x 4in) with a 3 mm smooth olive-shaped tip were positioned to target vascular channels in the liver for perfusion. The liver was first perfused an EGTA solution that comprised 0.14M NaCl, 50 mM KCL, 0.33 mM Na_2_HPO4, 0.44 mM KH_2_PO4, 10 mM Na-HEPES, 0.5 mM EGTA, 5 mM Glucose, and 4 mM NaHCO_3_ with pH adjusted to 7.2 using a Masterflex peristaltic pump (Cole-Palmer, Niles, IL, USA) set at a rate of 10 revolutions per minute (rpm), for approximately 1 h at 37 °C^32^.

Following perfusion with the EGTA buffer, the solution was replaced with a collagenase solution which comprises 0.1 % collagenase, 5 mM CaCl_2_, and 4 mM NaHCO_3_ in 1X HBSS solution (0.14 M NaCl, 50 mM KCL, 0.33 mM Na_2_HPO4, 0.44 mM KH_2_PO4, 10 mM Na-HEPES), pH 7.5. The perfusion was maintained at a rate of 8 rpm for approximately 1 h or until visible sign of digestion identified by liver indentation following gentle pressure or obvious cell dissociation through the Glisson’s capsule that overlays the liver. The digested liver is collected into a tissue culture dish containing chilled Gibco’s Williams media that is supplemented with 5% FBS, 1% glutamine, and antibiotics. Cell suspensions were filtered through sterile folded gauze, centrifuged at 50 g, 4°C for 5 min. The centrifugation step was repeated twice, and the resulting hepatocytes were assessed for viability using trypan blue dye. Cultures plates were coated with collagen (Collagen, Type 1 from rat tail, Sigma, Saint Louis, MO) diluted 1 to 50 ratio in sterile H_2_O prior to cell seeding. Hepatocytes were allowed to adhere overnight before further experiments.

### 2.7. Seahorse mitochondrial assay

To assess cellular respiration in primary baboon hepatocytes from control and MUN baboons, we used Agilent Seahorse XF96 Extracellular Flux Analyzer (North Billerica, MA, USA). Hepatocytes were plated in a collagen-coated 96-well seahorse plate at a density of 40,000 cells per well. The XFe96 sensor cartridges were hydrated overnight with H_2_O at 37 °C and replaced with seahorse XF calibrant 1 h before the assay. Oxygen consumption rate (OCR) and extracellular acidification rate (ECAR) were measured under basal condition and in response to serial injection of mitochondrial inhibitors including 1.5µM Oligomycin (to inhibit ATP synthase), 0.5 µM FCCP (Carbonyl cyanide-p-trifluoromethoxyphenylhydrazone; a mitochondrial uncoupler to measure maximum respiration) and 0.5 µM antimycin A and rotenone cocktail (to inhibit electron flow through the mitochondrial electron transport chain). We also determined OCR in response to 2 h exposure to 1 mM glucose or 100 µM H_2_O_2_ to model metabolic and oxidative stress respectively prior to the seahorse assay. OCR were normalized to cell density per well measured by a live-cell imaging system (IncuCyte S3, Santorius Corporation, Edgewood, NY, USA). Data were processed using the Agilent wave software.

### 2.8. Hepatocyte mitochondrial membrane potential

Mitochondrial membrane potential was determined using tetramethylrhodamine ethyl ester (TMRE) kit from Abcam. TMRE is a cell-permeant dye that accumulates in active mitochondria due to their relative negative charge. Hepatocytes (5,000 cells) were seeded in black-walled 96-well plates and incubated with media containing 200 nM TMRE for 20 min at 37 °C. Following incubation, cells were washed with phosphate-buffered saline, and fluorescence intensity captured using the IncuCyte (Red excitation: 567-607 nM and emission: 622-704 nM). FCCP (20 µM) was used as internal control as it prevents TMRE staining, and its signal was used to set the minimum intensity threshold for TMRE during data analysis. TMRE fluorescence intensity per image was normalized to phase image area.

### 2.9. Hepatocyte protein expression by immunoblotting

Cell homogenates for immunoblotting were prepared using commercially available RIPA buffer (Thermo Scientific, Waltham, MA, USA). The concentration of protein in the homogenates was determined by colorimetric protein assay^33^. Equal amounts of protein extract (15 µg) were separated by SDS-polyacrylamide gel electrophoresis (5 % staking and 12 % resolving gel) and transferred to a nitrocellulose membrane. Antibodies for individual components of the mitochondrial electron transport chain (ETC) complexes I-V (NDFUB8, SDHB, UQCRC2, MTCO1 and ATP5α), mitofusin 1 (MFN 1), dynamin-related protein 1 (Drp1), optic atrophy protein 1 (OPA1), peroxisome proliferator-activated receptor-gamma coactivator 1-alpha (PGC1a), catalase and superoxide dismutase 2 (SOD2), were incubated overnight in 2% BSA at 4°C. ETC complexes antibodies were assayed together as part of total OXPHOS antibody cocktail. Other primary antibody details are provided in table 1. Protein bands for each sample were visualized using LI-COR imaging system after 1 h incubation with LI-COR IRDye® 800CW goat anti-mouse and anti-rabbit secondary antibodies (LI-COR Biosciences, Lincoln, NE, USA). All immunoblots were quantified using LI-COR Image Studio Lite software.

**Table 1:** Primary antibodies for immunoblotting.

| Antibody | Dilution ratio | Company | Cat # |
| --- | --- | --- | --- |
| Total OXPHOS | 1:2,000 | Abcam | ab110411 |
| MFN1 | 1:1,000 | Abcam | ab221661 |
| Drp1 | 1:1,000 | Cell Signaling | 8570 |
| OPA1 | 1:1,000 | Cell Signaling | 80471 |
| PGC1a | 1:500 | Cell Signaling | 2173 |
| Catalase | 1:1,000 | Cell Signaling | 14097 |
| SOD2 | 1:1,000 | Abcam | ab13533 |
| Vinculin | 1: 10,000 | Cell Signaling | 4650 |

### 2.9. Statistical analysis

Data were analyzed by two-way analysis of variance (ANOVA) followed by Tukey post hoc test for multiple comparison and unpaired t-test when comparing effects between two groups. The ANOVA was weighted using SEM to account for variability in technical replicates. We did not observe any significant variation in hepatocyte data generated from the left and right liver lobe and were therefore pooled for subsequent analyses. Control male and female baboons were aged between 13.6-18.0 and 13.3-16.5 years, respectively, while MUN baboons were aged 13.7–16.2 years for males and 13.3-16.5 years for females. Data are presented as mean □±□ SEM; p<0.05 is considered statistically significant. All analyses were carried out using GraphPad prism 9.

## 3.0 Results

### 3.1. Liver metabolites

The concentrations of glycogen, choline and lipid in the liver measured by 1H-MRS were similar in both male and female MUN offspring when compared to their control counterparts. When male and female data were combined, the levels of these liver metabolites remained comparable between control and MUN baboons. Liver lipid concentration tended to be higher in control (p=0.095) relative to MUN baboons though these did not reach statistical significance (Fig. 1).

**Fig. 1:**
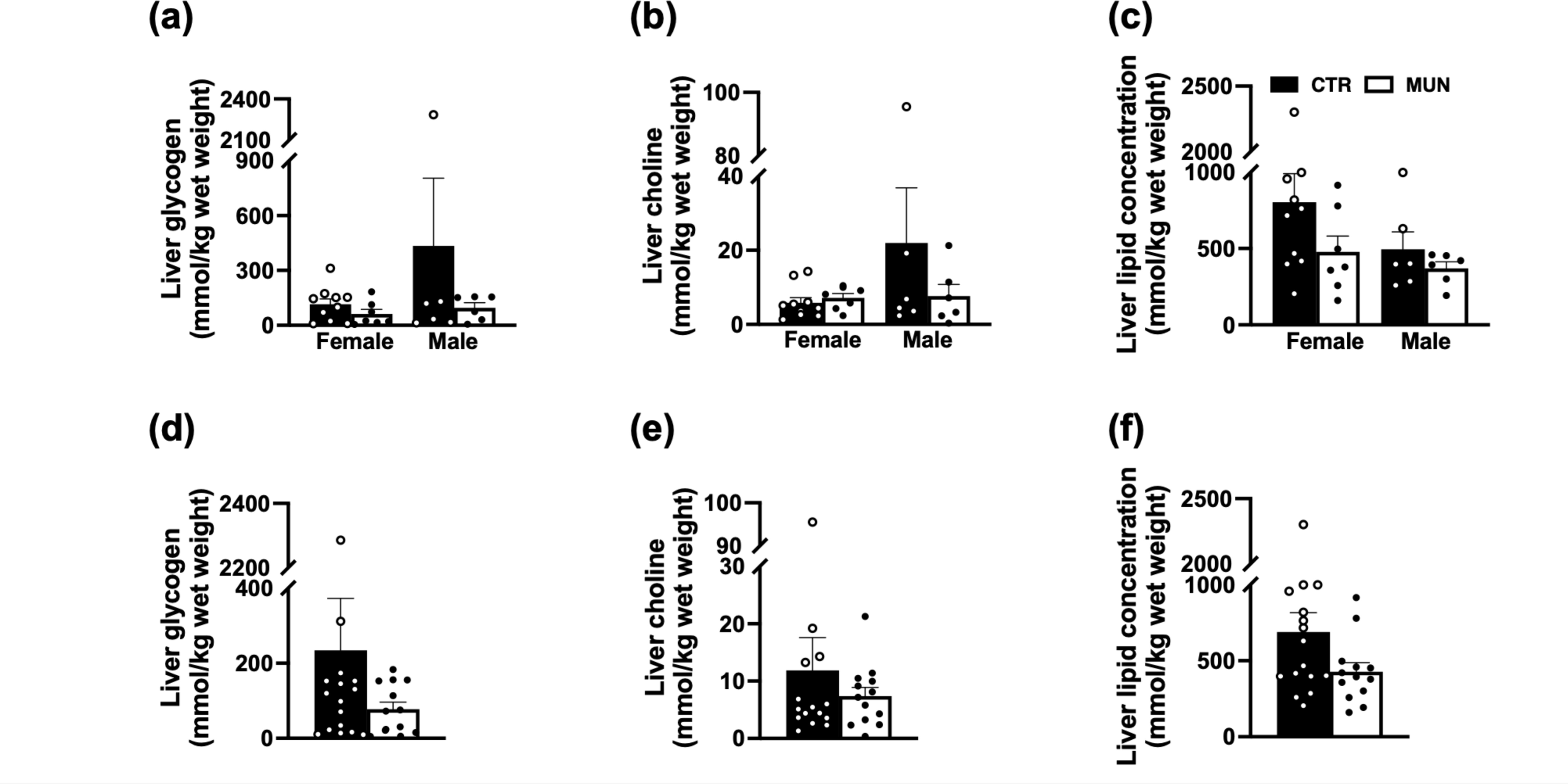
Liver metabolites in control and MUN baboon offspring. Liver metabolites were determined by magnetic resonance spectroscopy in female and male subjects from control and MUN groups (a) Liver glycogen concentration. (b) Liver choline concentration. (c) Liver lipid concentration. (d) Liver glycogen concentration for combined sexes (b) Liver choline concentration for combined sexes. (c) Liver lipid concentration for combined sexes. Black bars represent control baboons while clear bars are MUN baboons. Each dot represents an individual animal. Data are expressed as mean ± SEM, sample size and age range (control; n=10 for females, 12.4-16.4 years, 6 for males, 12.1-17.2, MUN; n=7 for females, 12.3-14.7 years, 6 for males, 12.4-15.0 years). For interaction of sex and treatment, data were analyzed by two-way ANOVA, while student’s t-test was used for treatment group comparison using GraphPad prism 9. Abbreviations: CTR; Control, MUN; maternal undernutrition.

### 3.2. Anthropometric variables and relative liver weight

Animals used in this study were aged-matched and there was no effect of MUN on body weight or body mass index (BMI) at this stage in later life. Overall, male baboon body weight was significantly higher than female in both control and MUN offspring. This sex-related difference in body weight is a well-recognized physiological factor. The relative liver weight, determined by the ratio of liver weight to body weight was similar between MUN and control offspring (Table 2).

**Table 2:**
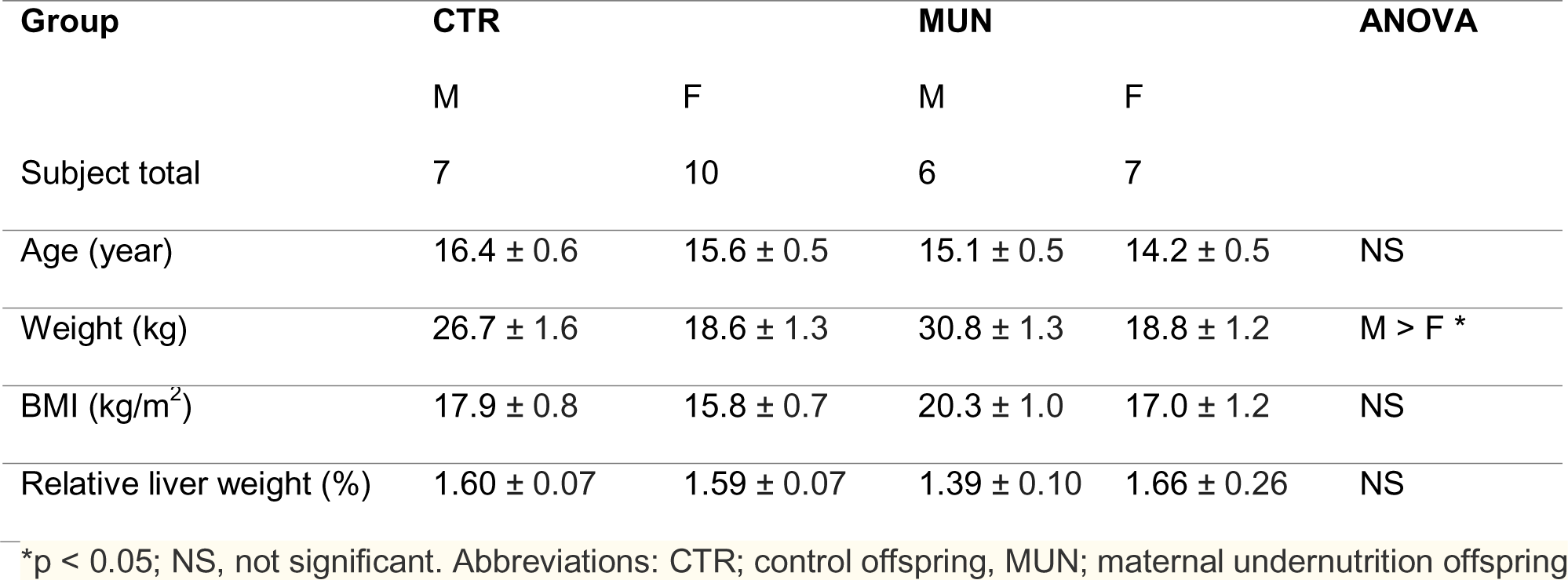
Anthropometric measures and relative liver weight in adult control and MUN baboons.

### 3.3. Liver enzymes

We compared plasma concentrations of AST, ALP, and ALT between control and MUN offspring as markers of liver function to determine the long-term impact of developmental undernutrition in adulthood. There were no significant differences in plasma liver enzyme levels between the groups when analyzed by sex independently. However, combined male and female data of control and MUN offspring showed significantly higher AST concentrations in MUN compared to control while ALP and ALT concentrations remained similar (Fig. 2).

**Fig. 2:**
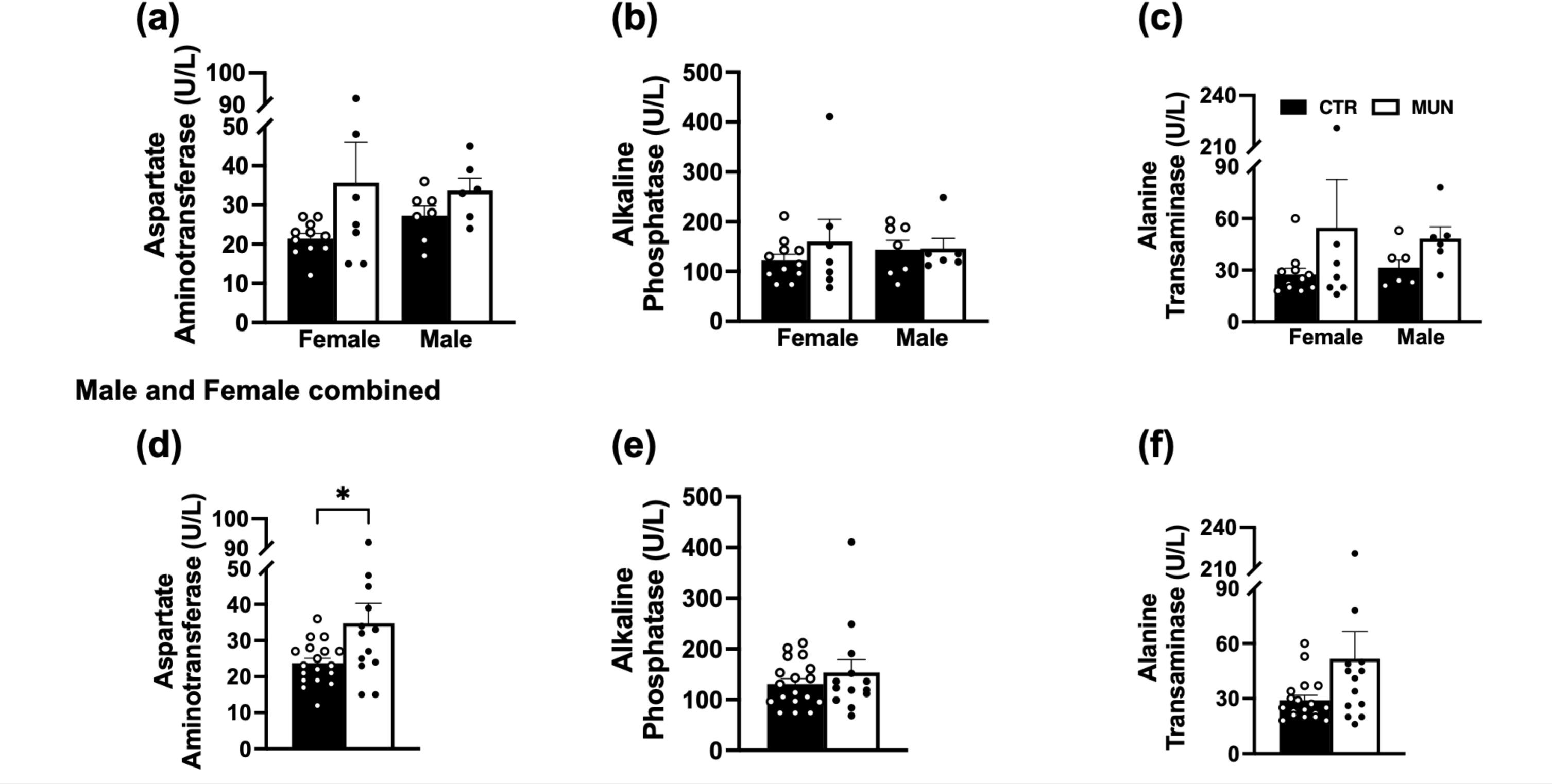
Plasma enzyme levels in control and MUN baboon offspring. (a) Aspartate aminotransferase levels in female and male subjects in the control and MUN groups. (b) Alkaline phosphatase levels in control and MUN baboons of both sexes. (c) Alanine transaminase levels in female and male baboons from control and MUN groups. (d) Aspartate aminotransferase levels for combined sexes in control and MUN groups. (e) Alkaline phosphatase levels for combined sexes in control and MUN groups. (f) Alanine transaminase levels for combined sexes in control and MUN groups. Black bars are for control baboons while clear bars are MUN baboons. Each dot represents an individual animal. Data are expressed as mean ± SEM, sample size (control; n=10 for females, 6 for males, MUN; n=7 for females, 6 for males). Age range: Control female and male baboons; 13.3-17.8 and 13.6-18.0 respectively, MUN females; 13.1-16.0 years, MUN males; 13.4–16.0 years. Two-way ANOVA was used determine sex and treatment interactions. When sexes were combined, difference between control and MUN baboons were determined by student’s t-test using GraphPad prism 9.

### 3.4. Hepatocyte number and viability

We next asked whether MUN had affects in adults at the hepatic cellular level using isolated hepatocytes from these animals to determine if MUN induce changes in cell physiology using readouts of cell viability and mitochondrial function (discussed below). Figure 3 shows that the number of live hepatocytes per gram of liver tissue as well as hepatocyte viability were similar between control and MUN baboons.

**Fig. 3:**
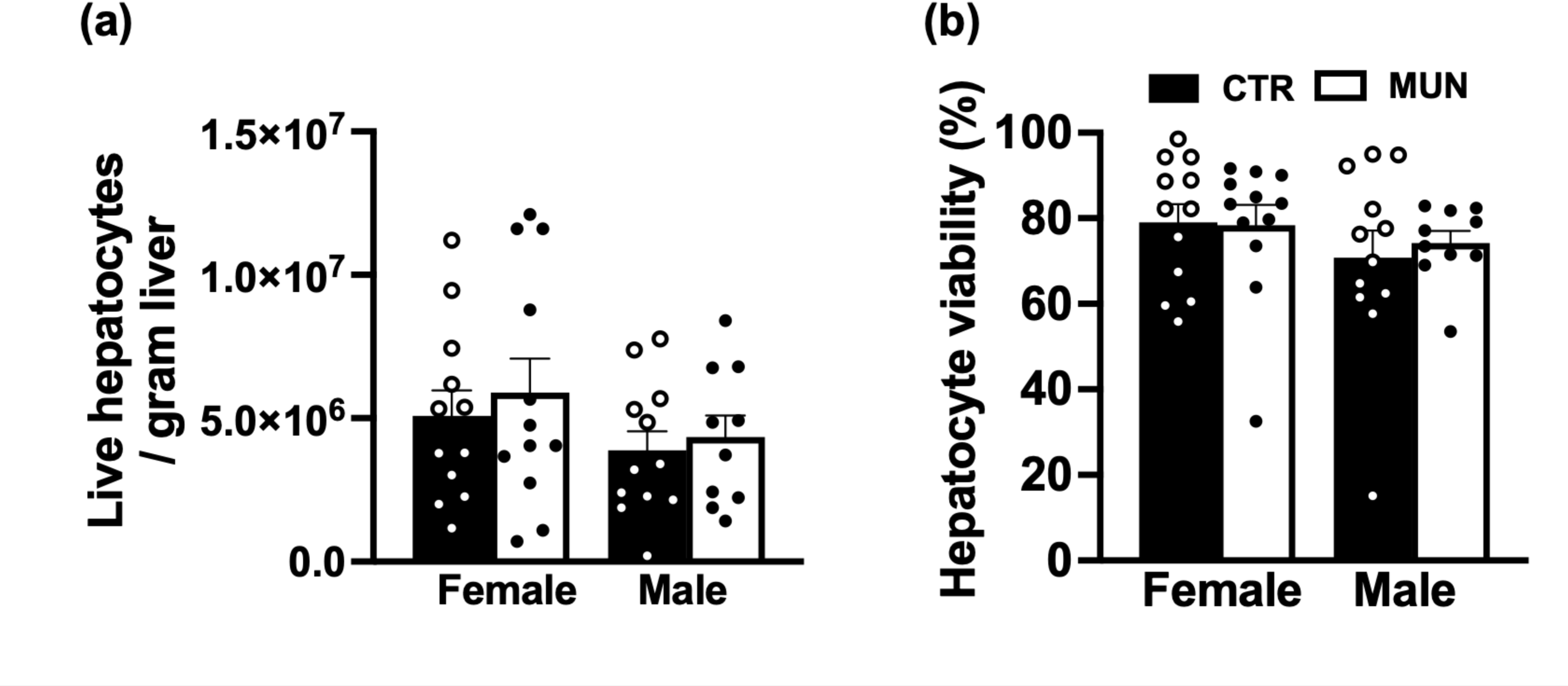
Viability of hepatocytes derived from control and MUN baboon offspring. (a) Number of live hepatocytes per gram of liver tissue (b) Hepatocyte viability. Data from left and right liver lobes were combined given that there were no lobe-specific differences. Each dot represents data point from individual liver lobes of each animal. Sample size; control females, n=6, control males, n=6, MUN females, n=6, MUN males, n=5.

### 3.5. Hepatocyte bioenergetics

We assessed hepatocyte bioenergetics using a mitochondrial stress test and show that OCR and ECAR were significantly higher in female-derived hepatocytes compared to males in control offspring. However, in MUN offspring there were no sex-differences in OCR and ECAR between male and female suggesting perhaps that early-life nutrient reduction may affect hepatic respiration sexual dimorphism. OCR parameters including basal respiration, ATP-linked respiration, maximal respiration, and spare respiratory capacity and ECAR were similar between in control and MUN female offspring. However, in male offspring, these parameters were significantly higher in hepatocytes from MUN offspring compared to hepatocytes from control animals (Fig. 4 a-d). Thus, MUN appears to have a male-specific effect on hepatic mitochondrial bioenergetics. The ratio of OCR to ECAR was also similar between the experimental groups (Fig. 4 e and f) suggesting no dramatic changes in mitochondrial fuel preference.

**Fig. 4:**
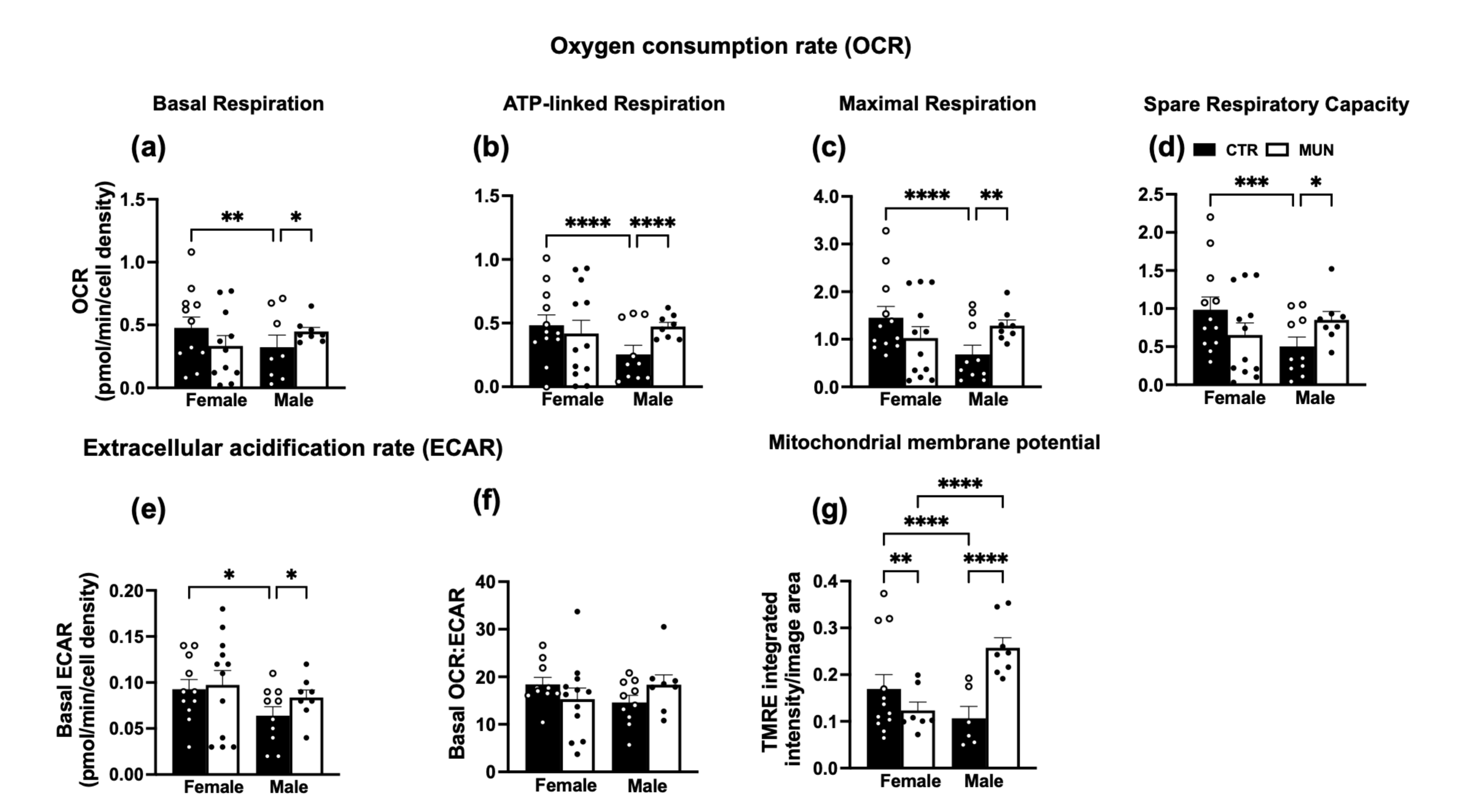
Cellular respiration in control and MUN baboon offspring. Oxygen consumption rate (OCR) and extracellular acidification rate in hepatocytes derived from MUN baboons and their control counterparts. OCR response to mitochondrial modulators such as oligomycin, FCCP and rotenone/antimycin were used to ATP-linked, and maximal respiration. Tetramethylrhodamine ethyl ester (TMRE) based assay was used to determine mitochondrial membrane potential (a) Basal respiration. (b) ATP-linked respiration. (c) Maximal respiration. (d) Spare respiratory capacity. (e) Basal ECAR. (f) Basal OCR to ECAR ratio. (g) Mitochondrial membrane potential. Data were expressed as mean ± SEM, with left and right liver hepatocyte data combined. OCR and ECAR were from 4 to 6 replicate samples and were measured using a seahorse XFe96 flux analyzer. OCR and ECAR data were normalized to cell density determined by a live-cell imager (IncuCyte). Seahorse assay sample size; control females, n=6, control males, n=5, MUN females, n=6, MUN males, n=4. TMRE assay sample size: control females, n=6, control males, n=3, MUN females, n=4, MUN males, n=4.

Mitochondrial membrane potential (MMP) was also significantly affected by MUN in isolated hepatocytes. Samples from MUN female offspring exhibit lower MMP when compared to their control counterparts, whereas in males, MMP was higher in MUN animals compared to controls. The MMP data correspond somewhat to the differences observed in OCR between MUN and control baboons, particularly in males. Additionally, we noted higher MMP in control females relative to males whereas the opposite was observed in MUN animals (Fig. 4 g).

### 3.6. Hepatocyte bioenergetics in response to low glucose and H_2_O_2_

We next asked whether MUN affected the response of hepatocytes from adult offspring to metabolic challenge in culture. In female-derived samples, hepatocytes exposed to low glucose (1mM) exhibited a significantly increased basal and ATP-linked respiration relative to standard glucose conditions in both control and MUN. However, maximum respiration and energy reserve were reduced in response to low glucose. These changes were observed in both control and MUN offspring suggesting MUN did not affect hepatocyte response to metabolic challenge in females (Fig. 5 a-d).

**Fig. 5:**
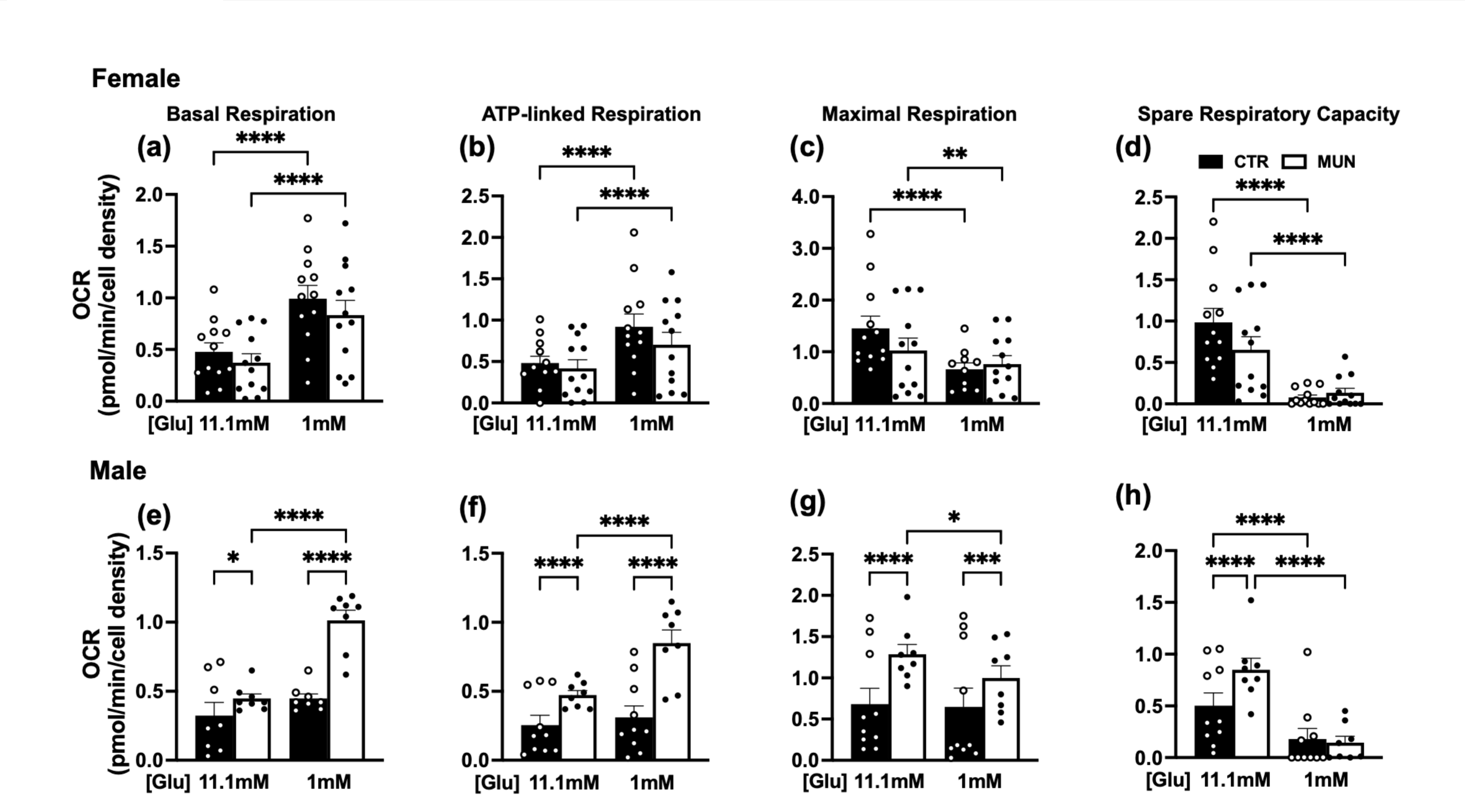
Cellular respiration in response to metabolic stress in control and MUN baboon offspring. Hepatocytes derived from male and female baboons in control and MUN groups were exposed to low glucose media (1 mM) for 2 h to model metabolic stress prior to mitochondrial stress test. Standard hepatocyte culture media contains 11.1 mM glucose. (a) Female basal respiration. (b) Female ATP-linked respiration. (c) Female maximal respiration. (d) Female spare respiratory capacity. (e) Male basal respiration. (f) Male ATP-linked respiration. (g) Male maximal respiration. (h) Male spare respiratory capacity. Data were expressed as mean ± SEM, with left and right liver hepatocyte data combined. OCR were from 4 to 6 replicate samples and were measured using a seahorse XFe96 flux analyzer. OCR and ECAR data were normalized to cell density. Sample size; control females, n=6, control males, n=5, MUN females, n=6, MUN males, n=4.

Unlike in females, male-derived hepatocytes showed low glucose-stimulated increase in basal and ATP-linked respiration only in MUN offspring. One interpretation is that increased OCR in MUN baboons might represent an increased energy demand in these cells under challenge. Hepatocytes from male MUN offspring also showed reduced maximal respiration following low glucose exposure. Low glucose challenge also ablated the difference in spare respiratory capacity we report between MUN and control hepatocytes under standard culture conditions (Fig. 5 e-f).

We also asked whether exposing hepatocytes to H_2_O_2_ (an inducer of oxidative stress) would reveal differences in bioenergetic response between control and MUN. However, this challenge did not significantly affect hepatocyte respiration except for a reduction in basal respiration in female MUN group following exposure to H_2_O_2_ (Supplemental Fig. 1). Furthermore, low glucose or 100 µM H_2_O_2_ elicited nearly identical changes in mitochondrial membrane potential similar to untreated cells in control and MUN baboons of both sexes (Supplemental Fig. 2).

### 3.7. Protein expression

We asked if the differences in bioenergetics in hepatocytes might be explained by mitochondrial content differences. The levels of OXPHOS proteins; complex I (NDUFB8), complex II (SDHB), complex III (UQCRC2), complex IV (MTCO1), and complex V (ATP5α) in isolated hepatocytes were not different between control and MUN offspring of both sexes (Fig. 7). This suggests the differences in bioenergetics we report is likely not related to mitochondrial abundance in these cell lines. Consistent with mitochondrial content, mitochondrial fusion proteins (MFN1 and OPA1) and fission protein (Drp1) were similar between the groups (Fig. 8 a-e). The marker of mitochondrial biogenesis (PGC1a) was also unchanged (Fig 8. f and h). The protein levels of antioxidant enzymes; catalase and SOD2 were not also altered in both male and female MUN baboons relative to their aged-matched controls.

**Fig. 6:**
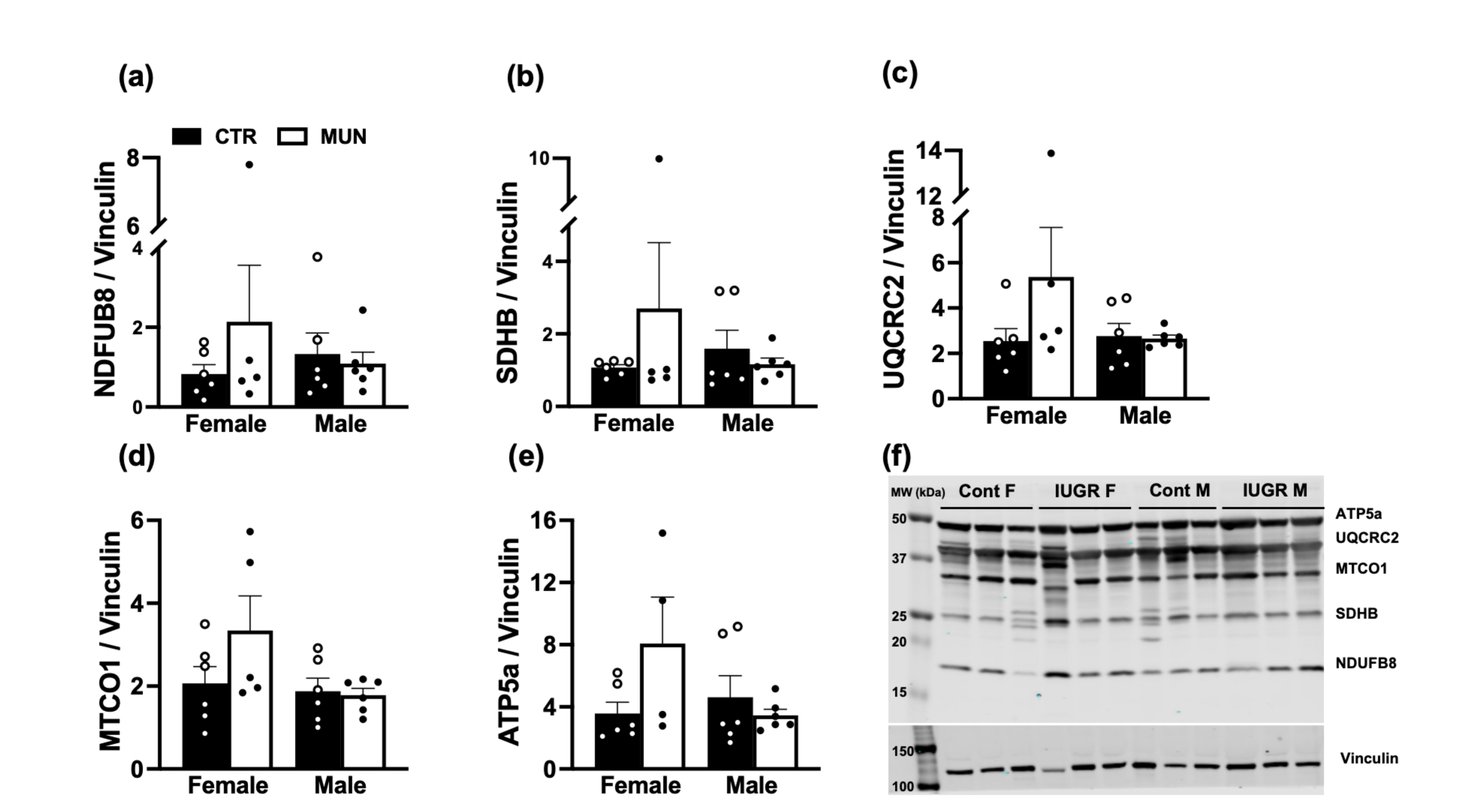
Mitochondrial electron transport chain subunits protein abundance in hepatocytes of control and MUN baboon offspring. Bar graphs present average normalized level of indicated mitochondrial subunit protein expression ± standard error of mean (SEM). Black bars represent control baboons while clear bars are MUN baboons. (a) Complex I (NDFUB8) protein expression (b) Complex II (SDHB) expression (c) Complex III (UQCRC2) protein expression (d) Complex IV,MTCO1 protein expression. (e) Complex V (ATP5a) protein expression (f) Representative photomicrograph of protein expressions in hepatocytes of control and MUN animals. Immunoblotting data are from left liver lobe hepatocyte. Sample size; control females, n=6, control males, n=5, MUN females, n=6, MUN males, n=5.

**Fig. 7:**
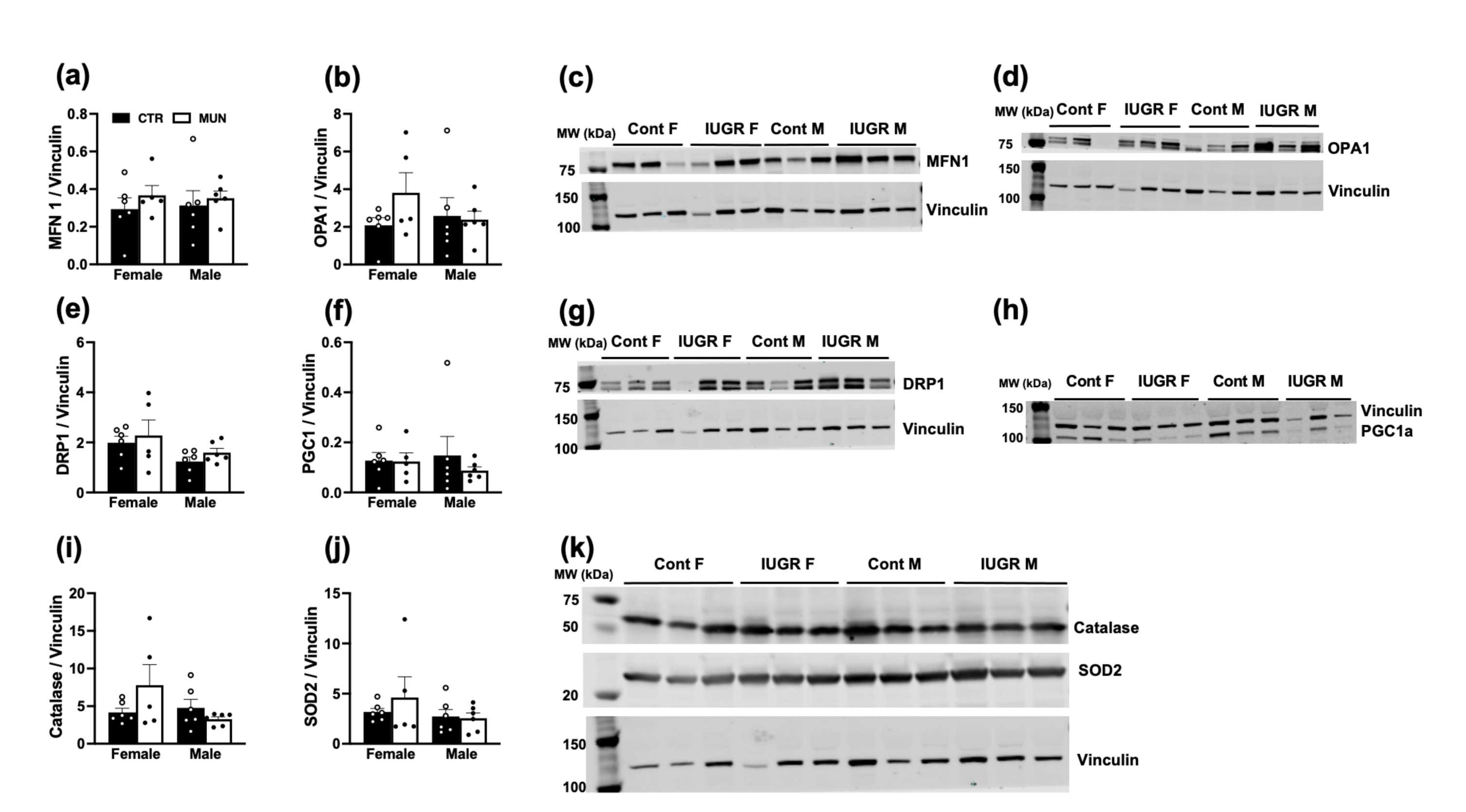
Levels of mitochondrial and antioxidant proteins in hepatocytes of control and MUN baboon offspring. Bar graphs present average normalized level of indicated mitochondrial subunit protein expression ± standard error of mean (SEM) determined by immunoblotting. (a) MFN I (b) OPA1 (c) Photomicrograph of MFN1 protein bands (d) Photomicrograph of OPA1 protein bands (e) Drp1 (f) PGC1a (g) Photomicrograph of Drp1 protein bands (h) Photomicrograph of PGC1a protein bands (i) Catalase (j) SOD2 (g) Photomicrograph of catalase and SOD2 protein bands. Immunoblotting data are from left liver lobe hepatocyte. Sample size; control females, n=6, control males, n=5, MUN females, n=6, MUN males, n=5.

## 4.0 Discussion

Our study shows that developmental programming imprints in baboons are notably evident at the cellular level, even in late adulthood, with primary hepatocytes derived from male MUN offspring highly sensitive to modulators of the mitochondrial electron transport chain, resulting in elevated OCR parameters relative to control offspring but without major changes to systemic liver function. The observed sexual dimorphism in mitochondrial OCR is consistent with the predominantly male-centric effect of programming in several metabolic studies. We previously reported male specific effects of MUN on genes regulating hepatic energy metabolism in fetal baboon liver and adipose tissue^34^. Postnatally, MUN also induces pericardial adiposity in 6-year-old male offspring but not in females^34^. Increased serum level of total cholesterol and low-density lipoprotein were also observed only in male MUN offspring at 9 years of age^35^. This study adds to the male-specific effects of MUN on mitochondrial bioenergetic parameters in hepatocytes derived from the aging offspring. Baboon lifespan has been reported as 11 or 21 years^36,37^ and thus, the animals used in this study aged between 13 and 18 years represent the transition period from mid-to late-life. Another interesting finding from this study is that mitochondrial OCR parameters are higher in female derived hepatocytes compared to male in control baboons, which agrees with other studies using liver tissue in mice^38^. It is not clear if this observation relates to estrogen signaling and translates to any protective effect in the female, in line with estrogen protective role against hepatic steatosis^39^. In male and female MUN offspring, we did not observe any sex differences, suggesting MUN abrogated this sexual dimorphism in mitochondrial respiration.

While most liver metabolic markers we examined were unaffected by MUN at the time of sampling, the changes in hepatocyte mitochondrial respiration may indicate a tendency for metabolic alterations particularly in the male animals. We view the similarity in liver metabolic phenotypes between MUN and control offspring as adaptive which could be altered in the presence of a second physiological insult overlaid on the perinatal exposure in adulthood. An adaptive response to poor perinatal nutrition triggers susceptibility to metabolic diseases in adulthood especially following exposures to nutritional challenge later in life. For example, postnatal catch-up growth following IUGR leads to obesity especially when the offspring are fed a hypercaloric diet^40^. Our study model did not incorporate any secondary challenge following the perinatal nutrient reduction but focused on the long-term perpetuation of programming effect, thus outcomes are representative of basal effects. However, our low glucose metabolic challenge results in hepatocytes are consistent with this idea.

We previously reported that MUN leads to reduced fetal liver weight, changes in fetal liver metabolites, smaller body weight at birth, juvenile prediabetic phenotype and altered lipid metabolism in young adulthood ^7,10,11,15,35^. There are few studies on the long-term impact of MUN on liver function in precocial species and to our knowledge, none specifically in aging NHP such as baboons, which share 96% genetic homology with humans^41^. Our study model could therefore address many confounding factors in human studies that limit clear delineation of the impact of MUN on liver function separate from secondary metabolic challenges like obesogenic feeding in adulthood. For example, in a Chinese famine cohort study, fetal and infant exposure to famine was linked to an increased risk of fatty liver disease in female offspring five decades later. However, these subjects were also obese, and the contribution of obesity in adulthood was not factored into the analyses^42^. Similarly, participants from the Helsinki Birth Cohort who were smaller during early childhood exhibited an elevated risk of fatty liver disease in adulthood but as obese subjects^43^. Thus, the relationship between early-life malnutrition and risk of hepatic disease, independent of adult overweight/obesity, remains unclear. In the present study, MUN offspring maintained similar body weight as control offspring, despite early postnatal catch-up growth^23^ and we did not see any clinical features of hepatic dysfunction in the MUN offspring; liver weight, hepatic glycogen, choline, and lipid content were similar to control offspring. Plasma markers of liver function were also unchanged except for higher AST levels in MUN offspring when male and female data were combined. Our results suggest that independent of an additional metabolic insult in adulthood, there are no signs of liver disease in aging MUN baboon offspring, despite potential cellular metabolic differences.

In a previous study by our group on the effects of MUN on liver function in aging sheep^44^, we observed that a 50 % nutrient restriction during early pregnancy did not alter liver weight in the female offspring at 6 years of age, average lifespan of a sheep is 7 years^45^, while liver glycogen content only tended to be greater in the MUN offspring compared to control. Meanwhile, the MUN offspring had elevated hepatic lipid levels and low expression of peroxisome proliferator-activated receptor-γ (PPARγ), a transcriptional regulator that modulates fat metabolism. Additionally, they had higher body weight compared to the control group^44^, suggesting a potential link between disturbance to liver metabolic function in MUN offspring and the occurrence of obesity. A different group demonstrated that MUN during early pregnancy in sheep resulted in small liver size in middle-aged male offspring, independent of changes to total body mass^12^. The changes in liver mass parallel a significant reduction in hepatocyte growth factor genes in the liver of the same animals. Even in the same species, there are variability in programming outcome on liver function likely due to the type of nutritional constraint, sex, body weight status and age at examination.

Early studies have demonstrated the hepatic mitochondrial dysfunction precedes the development of non-alcoholic fatty liver diseases^46^. Thus, animals can present normal circulatory metabolic phenotype concurrently with altered hepatic mitochondrial structure and function. We view the mitochondria as an early target organelle of developmental programming because the broad range of phenotypes of adverse perinatal exposures suggest the involvement of a common or integrative mechanism across different cells and tissues than individual molecular markers. Mitochondrial vulnerability to damage begins early. Mitochondria of fertilized oocytes are susceptible to damage from poor gestational conditions, these mitochondrial defects persist into fetal and postnatal life and are linked to increased risk of diseases including metabolic disorders^19,47–49^. We previously demonstrated that MUN during pregnancy increases activity of the fetal hypothalamic-pituitary-adrenal axis, evidenced by high cortisol and ACTH concentrations in near-term baboons^50^. Further, we have observed a rise in local cortisol production in male fetal liver under MUN conditions^34^. This hormonal milieu may contribute to programmed changes in liver mitochondria, potentially influencing hepatocyte bioenergetic capacity in adulthood, as demonstrated in this study.

Mitochondria participate in stress responses, with glucocorticoid receptors also expressed within the mitochondria^51^. In response to stress, cells consume more energy to maintain their viability, either for the synthesis of biomolecules required for growth or protective efficiency such as repair of cellular damage and return to homeostasis^52,53^. The higher OCR parameters in hepatocytes derived from male MUN offspring under standard cell culture conditions may reflect an adaptive response from perinatal nutritional exposure resulting in elevated energy demand to meet cellular processes such as gene transcription and translation which may be at a higher metabolic cost due to programming.

Alternatively, the higher OCR in male MUN hepatocytes may indicate a state of programmed hypermetabolism, that is, increased resting energy expenditure^51^ to maintain normal liver function following perinatal challenge. This energy budget may drive the similarity we observed in liver metabolites and markers of liver function at the systemic level between control and MUN offspring, albeit at a high energetic cost. The increased OCR is consistent with and corresponds to higher MMP; measured by accumulation of fluorescent cation TMRE within the mitochondrial matrix. The magnitude of MMP increase in male MUN offspring relative to control offspring suggests mitochondrial hyperpolarization likely due to hyperactivity of the proton pump in the mitochondrial respiratory chain. Mitochondrial hyperpolarization is associated with excessive ROS production which eventually triggers cell death^54^. It is important to note that changes in cell volume may influence TMRE fluorescence intensity. Since we normalized signals with cell area, changes in cell structure are unlikely to influence TMRE signals in our study. A sustained hypermetabolic state along with increased ROS production may eventually overwhelm defense capability of the antioxidant system, indicating potential for increased vulnerability to ‘wear-and-tear’ in the male MUN offspring. Moreso, stress-induced hypermetabolism has been reported to accelerate rates of aging^55^. Although, we did not measure ROS activity in the hepatocytes, the similar level of antioxidant proteins in both control and MUN offspring suggest the buffering capacity against oxidative stress may be lower in male MUN offspring. A related study also demonstrated that prenatal exposure to maternal stress is associated with higher leukocyte mitochondrial content and bioenergetic capacity in the offspring^56^, highlighting the potential significance of bioenergetic capacity among other mitochondrial phenotypes (e.g. mitochondrial biogenesis) in developmental programming. The similarity in protein markers of ETC subunits (complex I-V), mitochondrial biogenesis (PGC1a) and mitochondrial dynamics (MFN1, OPA1, and Drp1) between the experimental groups suggests that mitochondrial protein content or remodeling of mitochondrial network through fusion and fission processes does not contribute to the increased bioenergetic capacity of the male MUN offspring.

Similar to other reports showing increased cellular OCR in response to acute low glucose exposure^57^, basal and ATP-linked respiration in male MUN hepatocytes were further elevated by low glucose exposure, while control males were unaffected, which may indicate adaptation to metabolic challenge in control males. We can as well speculate that the high basal and ATP-linked OCR mimics a stress response to meet increased energy demand. It could also be related to oxidation of other fuel substrates like lipids in the absence of glucose, to sustain cell viability. The latter is more plausible since it has been demonstrated that low glucose increases dependency on fatty acid oxidation for basal mitochondrial metabolism^58^. Comparing this low glucose-stimulated increase in OCR parameters with the higher OCR in MUN male hepatocytes cultured in standard glucose media, fits the narrative that a similar mechanism that enhances basal respiration under low glucose raises hepatocyte respiration of male MUN offspring under standard culture conditions and thus represent stress responses. In females, low-glucose stimulated increase in basal and ATP-linked OCR were similar in control and MUN offspring, corroborating the perpetuation of male-specific effects in response to metabolic challenge. However, despite the stimulation of basal OCR by low glucose, energy reserve was drastically reduced following the metabolic stress, suggesting low glucose limits achievable maximal respiration. This reduction in energy reserve following exposure to low glucose is consistent with other studies^59^. In relation to oxidative stress challenge, we did not find major changes to the pattern of cellular respiration between the groups when hepatocytes were exposed to H_2_O_2_, which may relate to the concentration of H_2_O_2_ we tested.

In our baboon MUN model, aging male MUN offspring exhibit changes in mitochondrial bioenergetic parameters without concurrent systemic liver function alterations. This supports the idea that hepatic mitochondrial dysfunction may precede detectable circulatory defects^46^. There is a potential to detect hepatic defects if our study had more sample size as the combination of male and female data showed a significant increase in AST level in MUN offspring. Overall, this study suggests that changes to mitochondrial function may be an orchestrator of programming effect on liver function in adulthood.

## Acknowledgements

The authors express their gratitude to the Research Imaging Institute and the Southwest National Primate Center Staff for their continuous technical support in our baboon research program. We also acknowledge the administrative and technical support of Karen Moore, Wenbo Qi, Benjamin Morr, Kevin Thyne, Jonathan Gelfond, and Raechel Camones.

