## Supplementary material for "Perinatal maternal undernutrition in baboons modulates hepatic mitochondrial function but not metabolites in aging offspring": Supplentary Information

**Short title: Maternal undernutrition and offspring liver function**

**Daniel A. Adekunbi^1^, Bowen Yang^2^, Hillary F. Huber^3^, Angelica M. Riojas^2^, Alexander J. Moody^2^, Cun Li^4^, Michael Olivier^5^, Peter W. Nathanielsz^4^, Geoffery D. Clarke^2^, Laura A. Cox^5^, Adam B. Salmon^1,6^**

^1^Department of Molecular Medicine and Barshop Institute for Longevity and Aging Studies, The University of Texas Health Science Center at San Antonio, Texas, USA.

^2^Research Imaging Institute, Long School of Medicine, The University of Texas Health Science Center at San Antonio, Ant Texas, USA.

^3^Southwest National Primate Research Center, Texas Biomedical Research Institute, San Antonio, Texas, USA.

^4^Texas Pregnancy and Life-course Health Research Center, Department of Animal Science, University of Wyoming, Laramie, Wyoming, USA.

^5^Center for Precision Medicine, Department of Internal Medicine, Wake Forest University School of Medicine, Winston-Salem, North Carolina, USA.

^6^Geriatric Research Education and Clinical Center, Audie L. Murphy Hospital, Southwest Veterans Health Care System, San Antonio, Texas, USA.

**Corresponding author:** Dr. Adam Salmon, Barshop Institute for Longevity and Aging Studies, The University of Texas Health Science Center at San Antonio, 7703 Floyd Curl Dr, San Antonio TX, 78229. Texas, USA

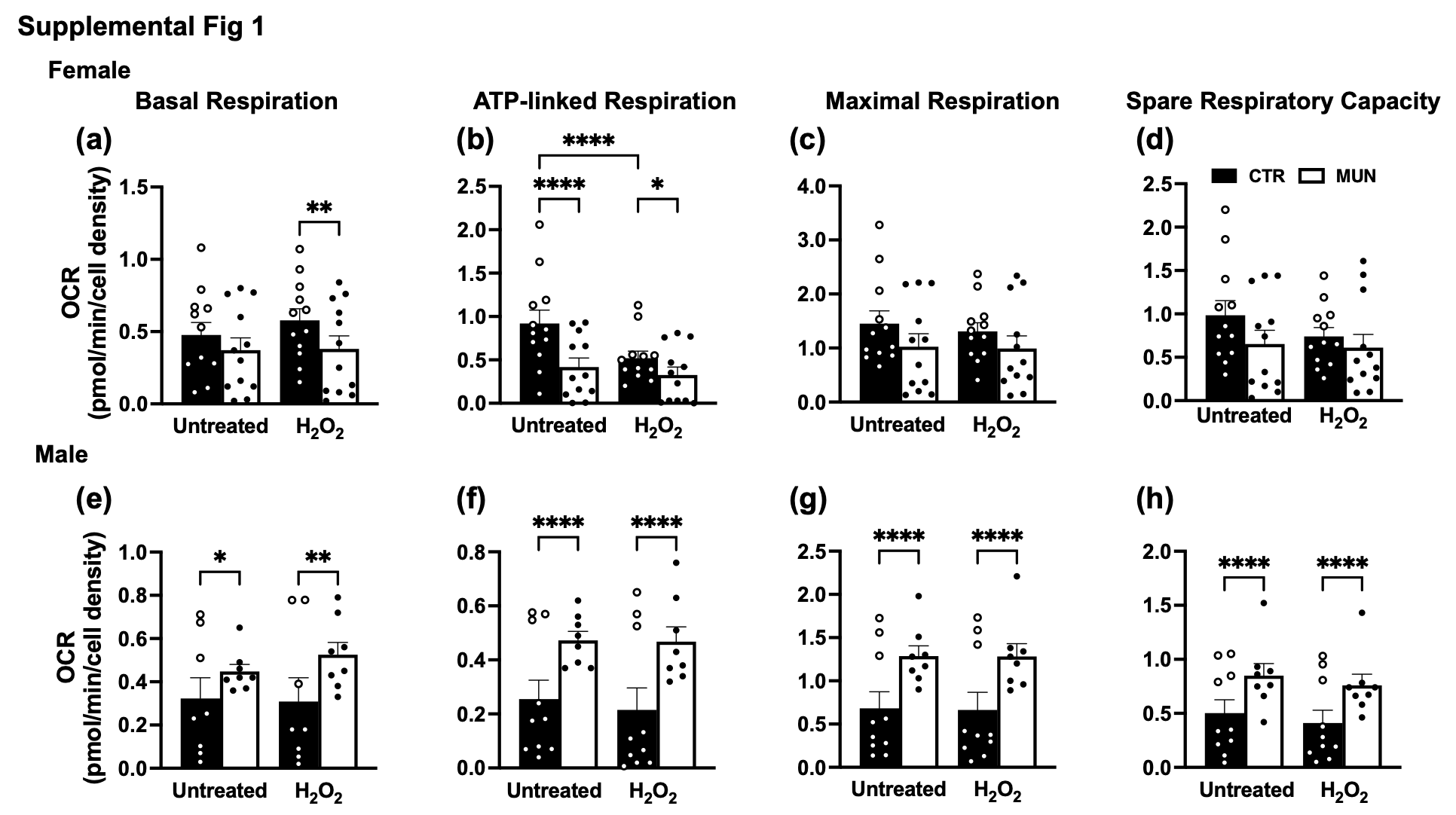

Supplemental Fig. 1: Cellular respiration in response to oxidative stress in control and MUN baboon offspring. Hepatocytes derived from male and female baboons in control and MUN groups were exposed to 100 µM H_2_O_2_ for 2 h to model oxidative stress prior to seahorse analysis. (a) Female basal respiration. (b) Female ATP-linked respiration. (c) Female maximal respiration. (d) Female spare respiratory capacity. (e) Male basal respiration. (f) Male ATP-linked respiration. (g) Male maximal respiration. (h) Male spare respiratory capacity. Data were expressed as mean ± SEM, with left and right liver hepatocyte data combined. OCR were from 4 to 6 replicate samples and measured using a seahorse XFe96 flux analyzer. OCR data were normalized to cell density. Age range and sample size: Control female and male baboons; control females, 13.3-17.8 years, n=6; control males, 13.6-18.0, n=5; MUN females, 13.1-16.0 years, n=6; MUN males, 13.4–16.0 years n=4.

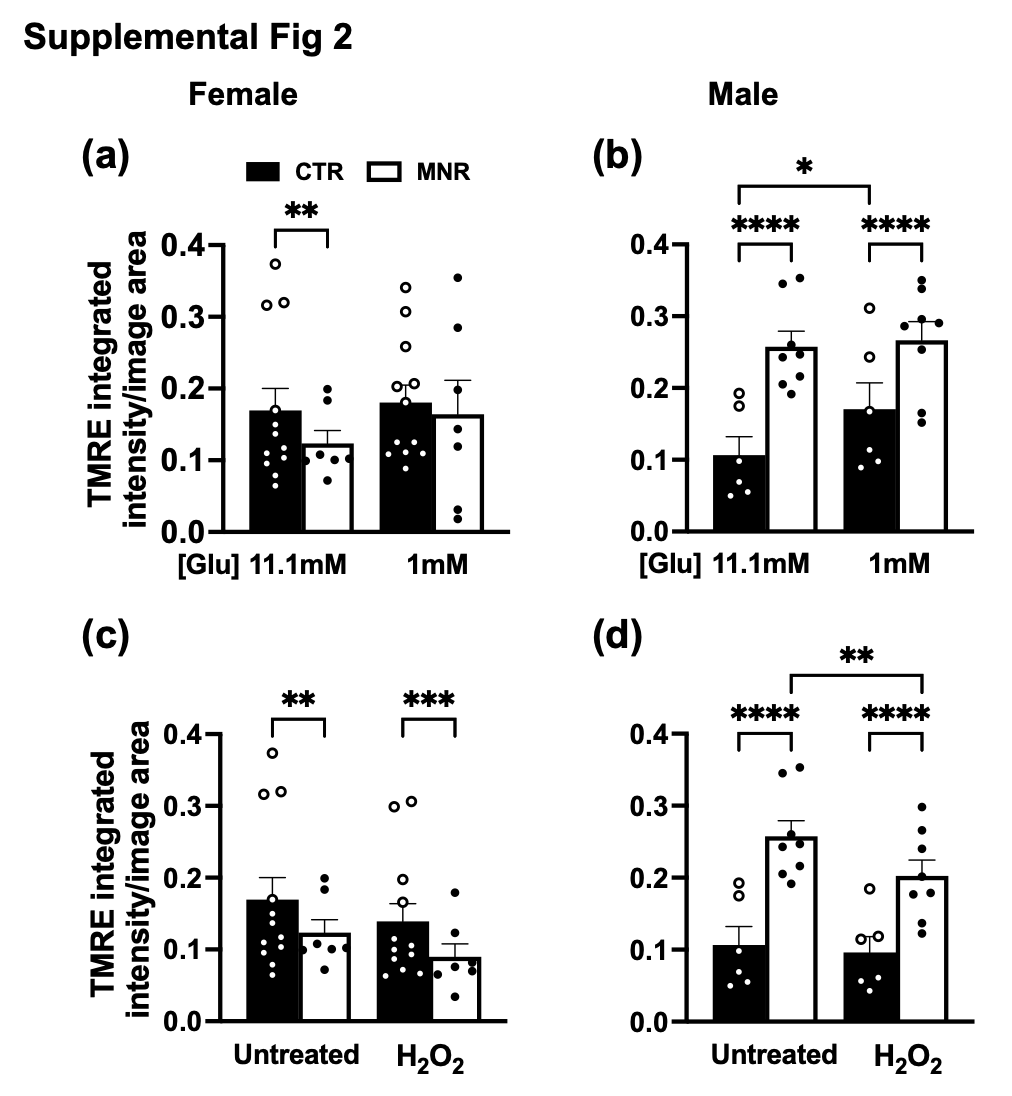

Supplemental Fig. 2: Mitochondrial membrane potential in response to metabolic and oxidative stress. Tetramethylrhodamine ethyl ester (TMRE) based assay was used to determine mitochondrial membrane potential (MMP) in control and MUN baboon hepatocytes following 2h exposure to 1 mM glucose or 100 µM H_2_O_2_ (a) MMP in female baboon hepatocytes in response to 1 mM glucose (b) MMP in male baboon hepatocytes in response to 1 mM glucose (c) MMP in female baboon hepatocytes in response to 100 µM H2O2 (d) MMP in male baboon hepatocytes in response to 100 µM H_2_O_2_. Data were expressed as mean ± SEM, with left and right liver hepatocyte data combined.
